# Mixed IgG Fc immune complexes exhibit blended binding profiles and refine FcR affinity estimates

**DOI:** 10.1101/2023.02.15.528730

**Authors:** Zhixin Cyrillus Tan, Anja Lux, Markus Biburger, Prabha Varghese, Stephen Lees, Falk Nimmerjahn, Aaron S. Meyer

## Abstract

Immunoglobulin (Ig)G antibodies coordinate immune effector responses by selectively binding to target antigens and then interacting with various effector cells via the Fcγ receptors. The Fc domain of IgG can promote or inhibit distinct effector responses across several different immune cell types through variation based on subclass and Fc domain glycosylation. Extensive characterization of these interactions has revealed how the inclusion of certain Fc subclasses or glycans results in distinct immune responses. During an immune response, however, IgG is produced with mixtures of Fc domain properties, so antigen-IgG immune complexes are likely to almost always be comprised of a combination of Fc forms. Whether and how this mixed composition influences immune effector responses has not been examined. Here, we measured Fcγ receptor binding to immune complexes of mixed Fc domain composition. We found that the binding properties of the mixed-composition immune complexes fell along a continuum between those of the corresponding pure cases. Binding quantitatively matched a mechanistic binding model, except for several low-affinity interactions mostly involving IgG2. We found that the affinities of these interactions are different than previously reported, and that the binding model could be used to provide refined estimates of these affinities. Finally, we demonstrated that the binding model can predict effector-cell elicited platelet depletion in humanized mice, with the model inferring the relevant effector cell populations. Contrary to the previous view in which IgG2 poorly engages with effector populations, we observe appreciable binding through avidity, but insufficient amounts to observe immune effector responses. Overall, this work demonstrates a quantitative framework for reasoning about effector response regulation arising from IgG of mixed Fc composition.

**Summary points:**

- The binding behavior of mixed Fc immune complexes is a blend of the binding properties for each constituent IgG species.
- An equilibrium, multivalent binding model can be generalized to incorporate immune complexes of mixed Fc composition.
- Particularly for low-affinity IgG-Fcγ receptor interactions, immune complexes provide better estimates of affinities.
- The FcγR binding model predicts effector-elicited cell clearance in humanized mice.

## Introduction

Antibodies are both a core component of adaptive immunity and a versatile platform for developing therapies. An antibody’s role in promoting immunity is defined by its selectivity toward a target antigen, as determined by its variable region, and its ability to elicit effector cell responses, defined by the composition of its constant, fragment crystallizable (Fc) region. Antibodies of the IgG type direct effector response by binding to the Fcy receptor (FcyR) family via their Fc-portion. FcγR activation is initiated through IgG-mediated clustering, which in turn is caused by the engagement of several antibodies on an antigen target forming an immune complex (IC). Depending upon the receptors included, this interaction may promote or prevent an effector response. This clustering mechanism ensures that more than one IgG is present whenever effector responses occur.

The immune response triggered by an IgG IC consisting of a specific Fc form, including subclass or glycosylation, is defined by its binding to specific Fcγ receptors, each of which differs in signaling effect and expression patterns^1^. Consequently, accurate estimates of IgG Fc-FcγR affinities are essential to understanding their effect. Most existing FcγR affinity measurements have been performed by surface plasmon resonance (SPR) using monovalent IgG^2,3^. SPR accurately assesses protein-protein binding kinetics, but many antibody-Fc receptor interactions are weak enough to fall outside the assay’s quantitative range when assessed in monovalent form. Clustering leads to avidity effects, wherein even weak interactions can cooperatively lead to strong binding^4^. Indeed, avidity is widely employed in natural and engineered systems to promote binding through low-affinity interactions^5^. Therefore, direct measurement of IC binding might more accurately quantify IgG Fc properties, particularly for low-affinity interactions. Measuring Fc binding as multivalent ICs additionally resembles the relevant *in vivo* context of effector responses^6^.

Physiological antibody responses universally involve Fc mixtures. For instance, during the course of infection, the composition of IgG subclasses shifts dynamically to different subclasses due to class switching^7^. Even when recombinantly manufacturing monoclonal therapeutic antibody preparations, heterogeneity exists in the glycosylation forms derived, and this glycan heterogeneity likely exists during endogenous antibody production as well^8,9^. With a mixture of antibodies of different Fc compositions but identical antigen binding, there might be an additive combination of effect, or a minor species (e.g., glycosylation variant) might present an outsized effect promoting or preventing effector responses. Therefore, knowledge of how these different forms influence the behavior of one another would allow one to modulate immune responses by adjusting the combination of different subclasses. With respect to therapeutic monoclonal antibody preparations, this would help guide the evaluation of biosimilars by determining whether glycosylation forms present at small fractions might influence overall therapeutic efficacy^10^.

After binding to Fc receptors, effector cell-elicited responses to IgG include several different functionally distinct mechanisms, including antibody-dependent cell cytotoxicity (ADCC) and phagocytosis (ADCP). Effector responses are coordinately regulated by the cell types present within a tissue^11,12^, the FcγRs expressed on those effector cells^13^, the Fc properties present within an immune complex^1^, and properties of antigen engagement^14,15^. Regulation at the Fc receptor and cell population level is a challenge to engineering antibodies with desirable cell-killing functions, as well as understanding both productive and pathogenic immune responses. Furthermore, it has become clear that in addition to NK cells, tissue-resident macrophages and bone marrow-derived monocytes participate in cytotoxic antibody-dependent target cell clearance. In contrast to NK cells (expressing only one activating FcγR, FcγRIIIA), these myeloid cell subsets express a broader set of activating FcγRs and the inhibitory FcγRIIB^13^. Thus, mixed IC may trigger all or specific subsets of activating/inhibitory FcγRs, resulting in further complexity. Despite the presence of and capacity to bind to multiple activating FcγRs on myeloid effector cells, our previous studies have demonstrated that individual IgG subclasses, such as mIgG2a/c for example, may mediate their activity through select activating FcγRs, indicating that there may be specialization in FcγR signaling^6^.

Our team recently demonstrated that a model of IC-FcγR binding accurately captured and could predict *in vitro* binding across various IgG isotypes^16^. Further, it could accurately predict antibody-elicited tumor cell killing in mice across antibodies of varied isotype, glycosylation status, and FcγR knockouts^16^. Directly quantifying and predicting cell clearance makes it possible to accurately anticipate and optimize for antibody-mediated therapeutic effects. However, it is still unclear whether such a modeling strategy can accurately predict the response of human immune cells, particularly given the divergent properties between the murine and human receptors^17–19^, and whether this modeling strategy can extend to ICs of mixed composition.

Here, we examined the binding properties of ICs with mixed IgG Fc composition. We quantified the binding of these ICs to each individual FcγR and observed that mixed-composition ICs resulted in a continuum of binding responses. A multivalent binding model extended to hetero-valent immune complex mixtures captured binding overall. However, surprisingly, it did not match certain low-affinity interactions^20^. Investigating the source of this discrepancy allowed us to improve the estimates of these interactions’ affinities. We additionally demonstrate that the binding model can be used to both predict *in vivo* effector responses in humanized mice and infer the cell types responsible for these responses. Thus, while antibody effector responses operate through a complex milieu of antibody species, Fc receptors, and cell types, IC profiling paired with modeling provides a framework to reason about the role of each molecular and cellular element.

## Results

### Profiling the binding effects of mixed-composition immune complexes

To determine the effect of having multiple Fc forms present within an immune complex (IC), we developed a controlled and simplified *in vitro* system. Like in previous work, we employed a panel of CHO cell lines expressing one of six individual human FcγRs^*16*^ (Fig. 1a). ICs were formed by immobilizing anti-2,4,6-trinitrophenol (TNP) human IgG on conjugates of TNP and bovine serum albumin (TNP-BSA) with an average valency of 4 or 33. IgG binding was then quantified after incubation with the cells, using a constant IC concentration of 1 nM (Fig. S1). In contrast to our previous work using a single IgG isotype, we assembled ICs from mixtures of each IgG isotype pair^*16*^. For each pair of IgGs, ICs were formed with a spectrum of six compositions of the IgG pair, including 100%/0%, 90%/10%, and 67%/33% mixtures. Combinations of 6 FcγRs, 2 valencies, 6 IgG pairs, and 6 IgG compositions resulted in 432 distinct experimental conditions. One-way ANOVA showed that more than 70% variance in the data are between experimental conditions rather than within them, indicating that more than 70% of the variance could be explained by biological differences. This suggests that, within each condition, measurements were consistent (Tbl. S1).

**Figure 1:**
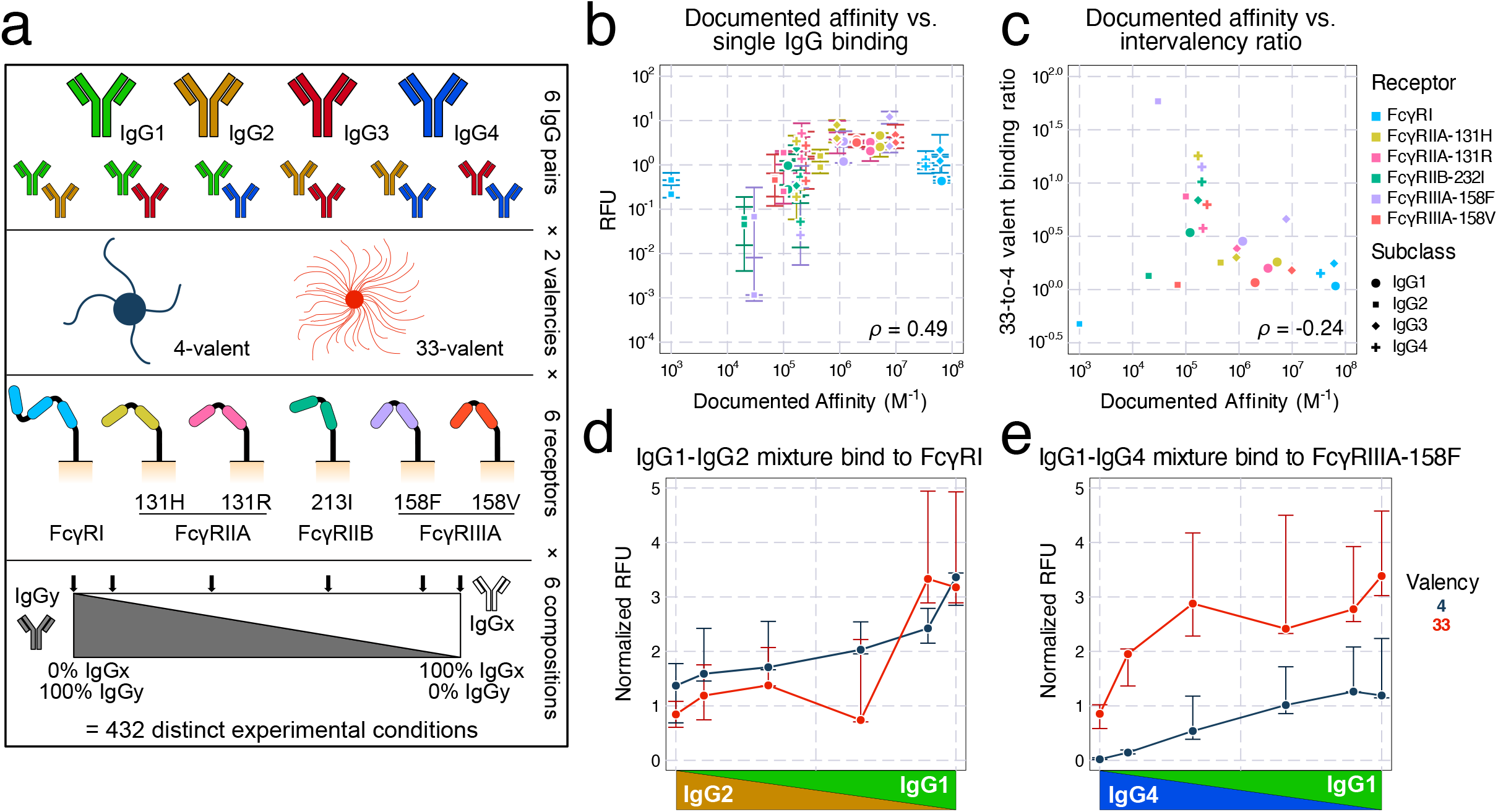
Profiling the binding effects of mixed-composition immune complexes. a) Schematic of the immune complex (IC) binding experiment. Individual or mixtures of IgG subclasses are immobilized on multivalent TNP complexes. The binding of these complexes to CHO cells expressing a single Fc receptor is then quantified. b) Measured binding in relative fluorescence units (RFU) versus the previously reported affinity of each interaction. Only single subclass conditions are plotted. Error bars represent the interquartile range of the measurements. c) The ratio of median binding quantified between valency 33 and 4, versus the reported affinity of the interaction. d) IgG1-IgG2 mixture binding to FcγRI shows appreciable binding, even though IgG2-FcγRI is documented to be non-binding. The RFU level here was normalized to match the FcγRI expression to the FcyRIIIA-158F expression (shown in e) on CHO cells. e) IgG1-IgG4 mixture binding to FcyRIIIA-158F.

Inspection of the resulting binding data revealed several expected patterns. Among the conditions with only one IgG present, the measured binding showed a strong, positive correlation with the documented IgG-FcγR interaction affinities (Fig. 1b). The higher valency ICs universally showed greater binding signal compared to their matching lower-valency counterparts, and there is an obvious negative trend between documented affinities and the ratio between the 33-valent and 4-valent complex binding (Fig. 1c). This trend is expected since, although complexes of both valencies can bind densely with high-affinity units, only high-valent complexes compensate for low affinity through avidity^*4*^. Therefore, while high-affinity complexes result in greater binding, low-affinity complexes have greater intervalency binding ratios. Finally, mixtures spanning 100% of one IgG isotype to another generally showed a monotonic shift with composition (Fig. S1). These patterns, along with their reproducibility (Tbl. S1), gave us confidence in the quality of the binding measurements.

We also observed several unexpected trends among the binding measurements. There was appreciable binding from IgG2-FcγRI interactions, despite this combination being reported as non-binding^*3*^ (Fig. 1d). We also saw an increase in binding along the shift from IgG4 to IgG1 with FcγRIIIA-158F, even though these two isotypes are documented to have identical affinity^*3*^ (Fig. 1e). These two observations are consistent with previous binding measurements using the same TNP-based IC system^*16*^.

To better visualize the binding measurements, we performed principal component analysis (PCA) on the median measurement of each condition, with each isotype mixture and valency as a sample and each receptor as a feature. The first principal component (PC1) explains more than 86% of the variance, and the first two components (PC1 and PC2) explain 93% (Fig. 2a). Inspecting the scores, we found that the 33-valent measurements are more broadly distributed, consistent with their greater expected binding (Fig. 2b/c). PC1 mostly separates IgG3 binding from other isotypes, reflecting that IgG3 has the greatest binding among IgG subclasses (Fig. S1). PC2 separated the genotype variants of FcγRIIA and FcγRIIIA and associated most strongly with IgG3 and IgG4 (Fig. 2d), reflecting that these two subclasses showed larger differences in binding with genotype (Fig. S1).

**Figure 2:**
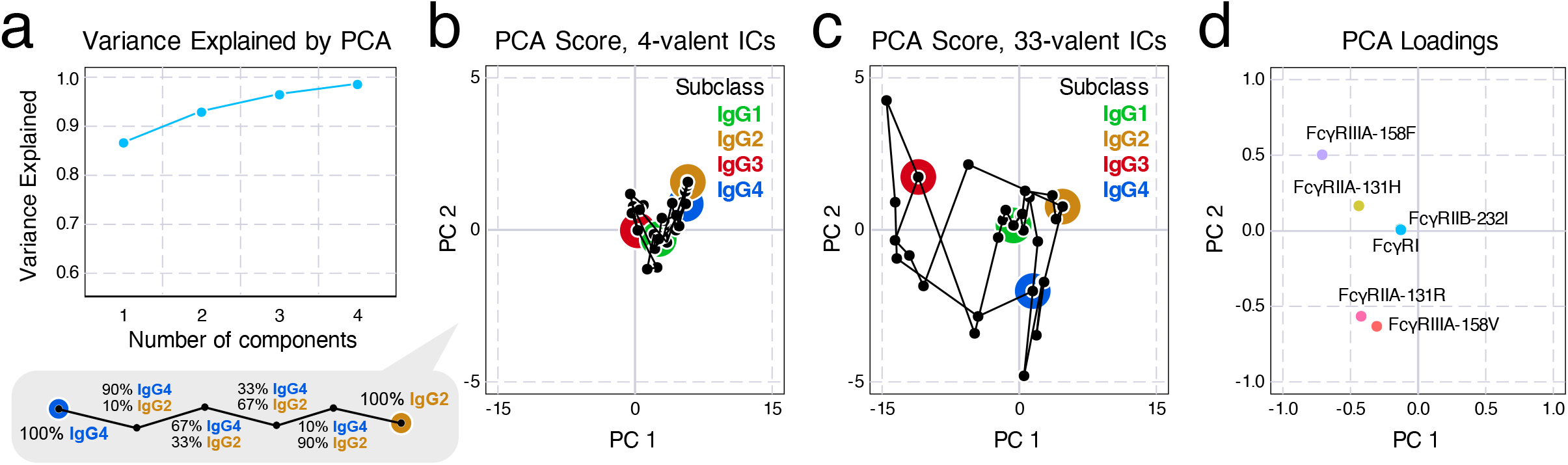
Principal component analysis (PCA) visualizes the variance in mixture binding measurements and their associated factors. a) Variance explained by each component in PCA. We found that the first two principal components (PC), PC1 and PC2, can explain more than 93% of the variance in the measurement. b,c) Scores of PC1-PC2 for immune complexes of average valency 4 (b) and 33 (c); PC1 shows that most of the variance in data comes from 33-valent complexes, especially in IgG3. d) Loadings of PC1-PC2 indicate that PC2 is mostly associated with separating the allelic variants of FcyRIIA and IIIA. FcyRI and FcyRIIb points overlap.

In all, these data support that TNP-assembled ICs provide a controlled *in vitro* system in which we can profile the effects of mixed IC composition on binding to effector cell populations. Quantifying binding using ICs may, in fact, provide more precise quantification of IgG-FcγR interaction affinities, particularly for lower affinity pairs, and mixed Fc composition ICs showed binding between that of the corresponding single Fc cases.

### A multivalent binding model accurately predicts *in vitro* IgG mixtures binding and updates Fc-FcγR affinities

To model the effects of polyclonal antibody responses, we extended a simple, equilibrium binding model that we have previously used to model antibody effector response^16,20^. Briefly, immune complexes are assumed to bind to FcγRs on the cell surface with monovalent binding kinetics, and then can engage additional receptors with a propensity proportional to their affinity (Fig. 3a). Though additional assumptions are not required for modeling ICs of mixed isotype composition, this extension leads to a large combinatorial expansion in the number of binding configurations. Through some properties of combinatorics, we derived simplified expressions for many macroscopic quantities to allow this model to scale to multi-ligand, multi-receptor, and multivalent situations^20^.

**Figure 3:**
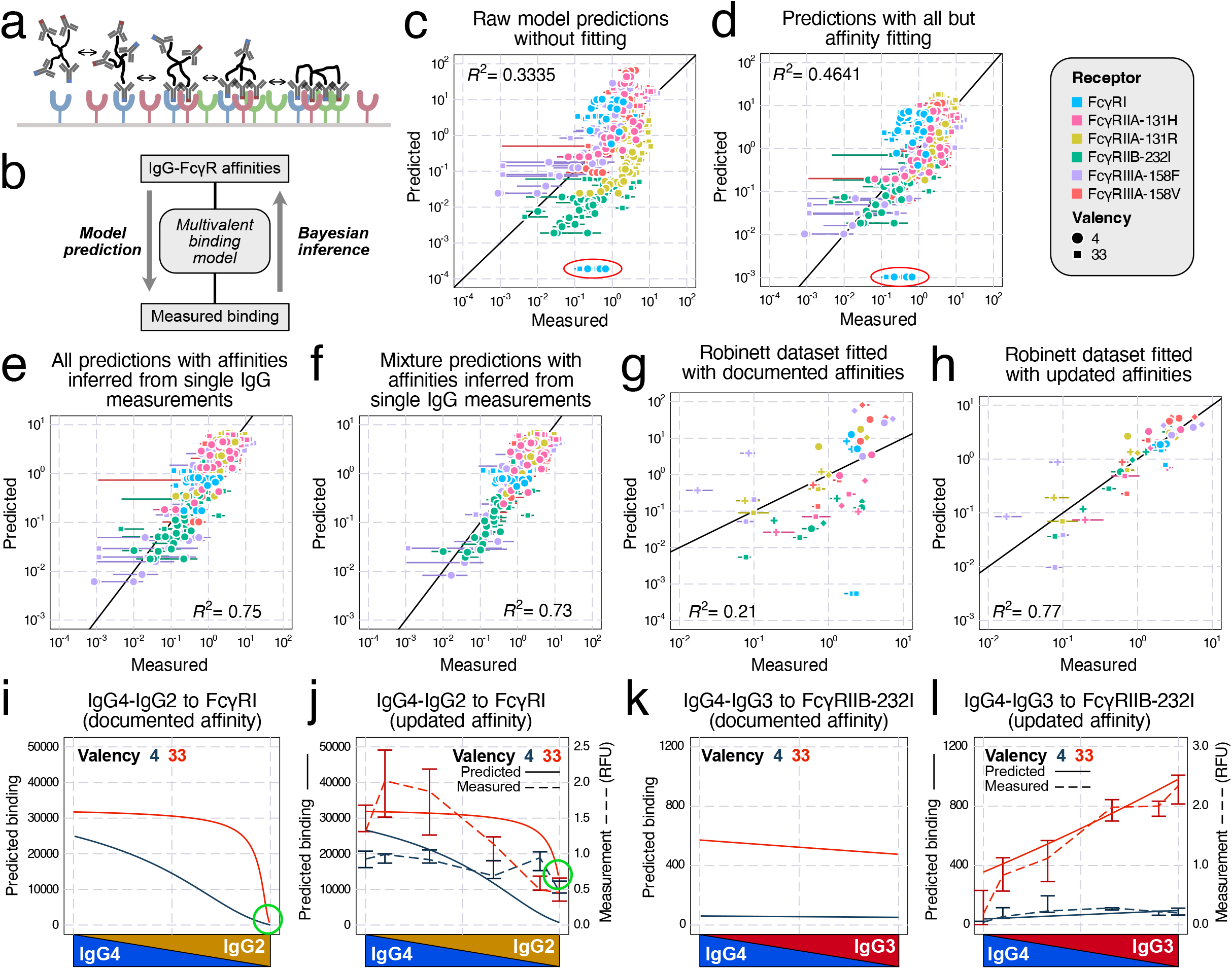
A multivalent binding model accurately accounts for *in vitro* binding of IgG mixtures. a) Schematic of the multivalent binding model. b) Schematic of the process of predicting binding with documented affinities and inferring affinities from measurements. c) Measured versus predicted binding by the binding model with raw parameters. Points also vary in the IgG subclass used, which is not indicated. d) Binding model prediction with all parameters but affinities fitted by Monte Carlo Markov Chain (MCMC). In (c) and (d), the IgG2-FcγRI outliers were circled in red. Since this interaction was previously reported as nonbinding, the actual predictions were all 0, only clipped to a nonzero value (1/10 of the next smallest value) to be plotted on logscale. e) Binding model prediction of all measurements (single and mixed IgG) with affinity inferred from single IgG measurements. f) Binding model prediction of mixture IgG measurements with affinities updated with single IgG measurements. g,h) Validate the updated affinities with a separate dataset^*16*^ by predicting the binding with either documented (g) or updated (h) affinities. i-l) Predicted binding of IgG4-IgG2 mixture to FcγRI (i,j) and IgG4-IgG3 mixture to FcγRIIB-232I (k,l), with either documented (i,k) or updated affinities (j,l, solid line and left axis) compared with measured binding (j,l, dashed line and right axis).

We first used the measured receptor expression (Tbl. S2) and documented affinities^3^ with the model and obtained reasonable agreement overall (Fig. 3c). While the predicted values mostly agreed with the measurements, there were several notable outliers, most prominently an underestimate of IgG2-FcγRI binding (Fig. 3c, red circle). To improve the measurement fit, we reversed the estimation process and used the measured binding to infer the interaction affinities via Markov chain Monte Carlo (MCMC) (Fig. 3b). We first created a baseline fit quality by fitting all but the affinities (e.g., receptor abundance and the crosslinking parameter 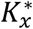, Fig. 3d). Although the fit improved, outliers persisted (circled in red in Fig. 3d). Therefore, we next performed the fitting while allowing the Fc-FcR affinities to vary. Although we only used the single-IgG measurements to infer the Fc affinities (Fig. 3e), we obtained much more accurate predictions for all measurements of both single and mixed IgG compositions (Fig. 3f).

To further confirm the generality of these updated affinities, we validated these new affinity estimates with an independent dataset collected in a previous study^16^. This previous study independently measured the binding of BSA-TNP complexes *in vitro* with two distinct average valencies (4 and 26), but only the binding of single IgG isotypes. We set the Fc affinities to either documented or updated values and let MCMC fit the other parameters. The new affinities resulted in a vastly improved agreement with the data (Fig. 3g/h).

To illustrate the impact of the affinity changes, we compared the binding predictions with two sets of affinities (Fig. S2/S3) to their corresponding measurements (Fig. S1). For FcγRI binding to IgG2-IgG4 mixtures, the experiment indicated that there is still notable binding with mostly or 100% IgG2, while IgG2-FcγRI was documented as non-binding^3^. The updated values amended the prediction and reflected this interaction, especially for the 33-valent complex (Fig. 3i/j, green circle). For FcγRIIB-232I binding to IgG3-IgG4, the documented affinities indicated there should be more binding to IgG4 compared to IgG3, contrary to our observation (Fig. 3k). The updated affinities instead accurately predicted the binding of all mixtures at both valencies (Fig. 3l). These examples demonstrate that the affinity adjustments greatly improved agreement with the binding measurements.

As our Fc affinity inference was constructed in a Bayesian fashion, both the prior (documented) and the posterior (updated) affinity values are represented as distributions accounting for uncertainty. Inspecting these updated distributions (Fig. 4a–d; Tbl. S3), we noted several trends. The model made the largest adjustments to the Fc affinities of IgG2 (Fig. 4b), followed by IgG4 (Fig. 4d). Most IgG1 (Fig. 4a) and IgG3 (Fig. 4c) affinities remained unmodified, except for a slight increase in their FcγRIIB-232I affinities. The most notable update occurred to IgG2-FcγRI. Previously reported as nonbinding, FcγRI was revised to be the highest affinity receptor for IgG2, consistent with the receptor’s high affinity to other human IgG subclasses. This discrepancy was reflected in the model prediction before affinity fitting, where the IgG2-FcγRI binding was the striking outlier (Fig. 3d/g). Another significant adjustment occurred with IgG3-FcγRIIB-232I. Although FcγRIIB-232I has a low affinity for all IgG subclasses, our update led to IgG3 being the strongest-binding subclass (Fig. 4c, S2 & S3). More subtle differences can be observed from specific model predictions (Fig. S2 & S3). The revised affinities showed a similar overall correlation with binding overall (Fig. 4e). The inter-valency binding ratios show a more prominent negative correlation, however, due to the movement of the IgG2-FcγRI outlier (Fig. 4f).

**Figure 4:**
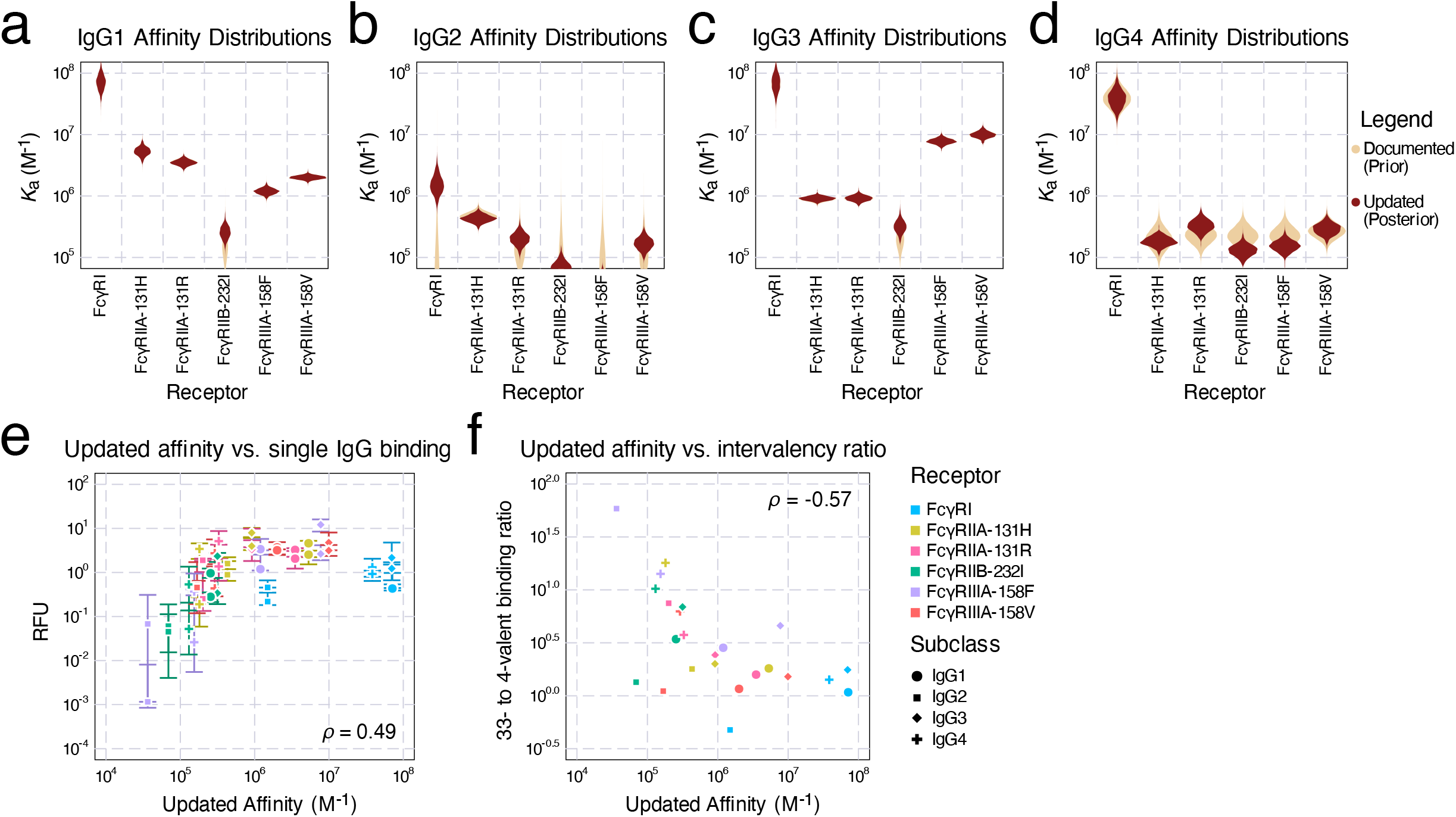
Inferred affinities from the binding data. a-d) The prior (documented) distributions of binding affinities (assume all follow log-normal distributions) and posterior (updated) affinities of IgG1 (a), IgG2 (b), IgG3 (c), and IgG4 (d). e) Updated affinities plot against the binding measurements of single IgGs. Error bars represent the interquartile range of the measurements. f) Updated affinities plot against the ratio of median binding between valency 33 and 4 complexes.

### Multivalent binding predicts antibody-elicited effector responses in humanized mice

We next sought to link the binding of ICs to their effects on the clearance of antigen targets *in vivo.* To quantify the antibody-driven activity of each effector cell, we first measured the binding of each human IgG subclass to immune effector cells *in vitro* in IC of two valencies, 4 and 33 (Fig. 5a–d). The measurements show that the binding amounts of IgG1 and IgG3 were generally about 10-fold higher in magnitude than those of IgG2 and IgG4. For the latter two subclasses, their 4-valent complex binding was almost negligible. In all cases except IgG2, neutrophils had more binding than classical and nonclassical monocytes.

**Figure 5:**
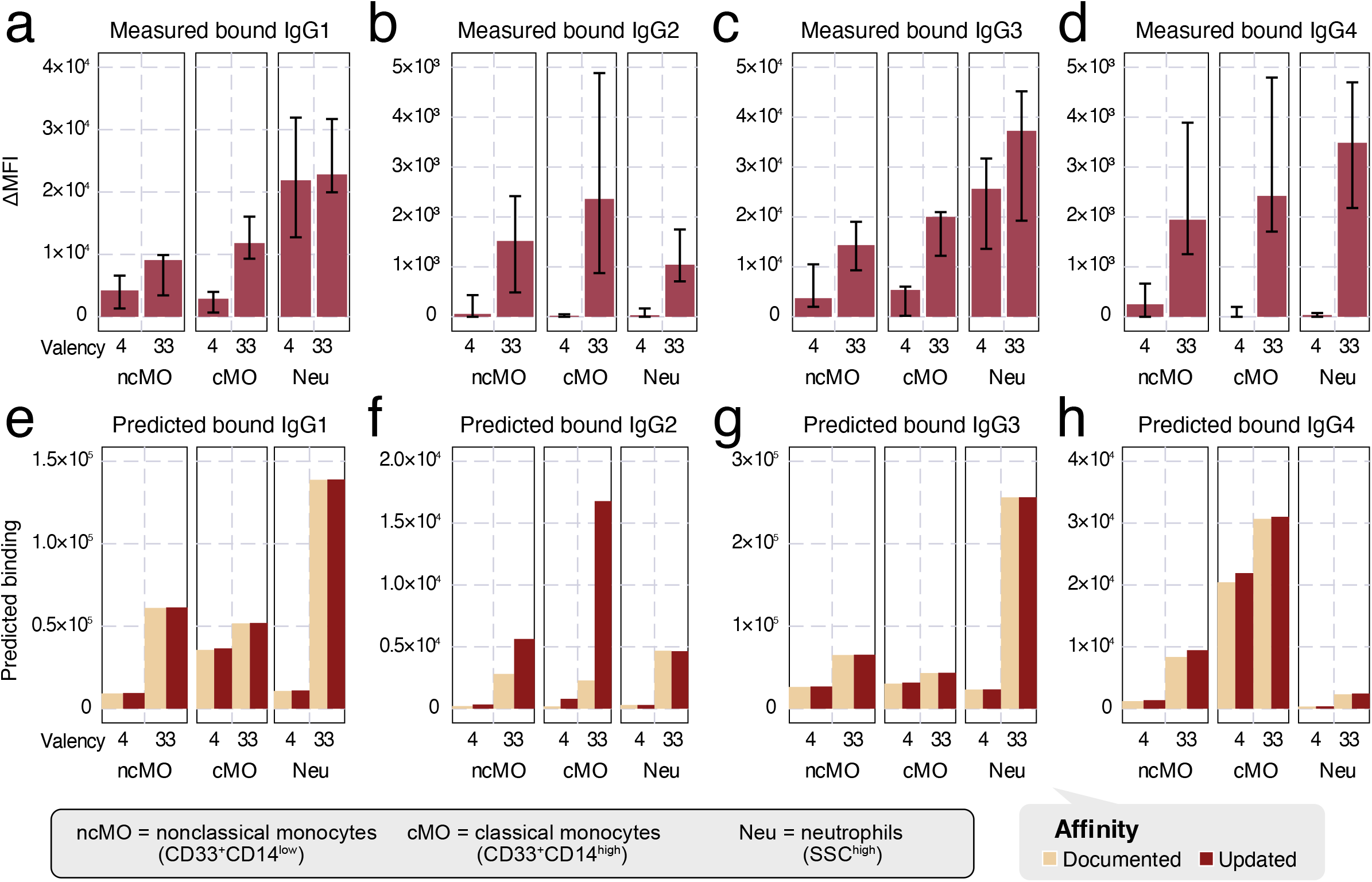
Predict IgG effector cell binding with the multivalent binding model. a-d) Measured *in vitro* binding of IgG1 (a), IgG2 (b), IgG3 (c), and IgG4 (d) IC of either 4- or 33- valent to selective immune effector cells, classical (cMO) or nonclassical (ncMO) monocytes and neutrophils (Neu). e-h) Model-predicted IgG1 (e), IgG2 (f), IgG3 (g), and IgG4 (h) IC of 4- or 33- valent binding on each effector cell type under documented versus updated affinities.

We predicted the same quantities of IC binding by the multivalent binding model with either the previously documented^3^ or updated affinities (Tbl. S4), and the quantification of FcγR abundance^13^ (Tbl. S6, Fig. 5e–h). These estimated binding amounts broadly aligned with the measurements. Between the two sets of affinities, the predictions for IgG1 and IgG3 remained almost identical (Fig. 5e/g), while some differences were reflected in IgG2 and IgG4 (Fig. 5f/h), consistent with the affinities changing more for IgG2 and IgG4 (Fig. 4a–d). Most prominently, the predicted binding to nonclassical and classical monocytes was adjusted to be much higher for 33-valent IgG2 (Fig. 5f), better matching the measured values (Fig. 5b). These changes indicate that the updated affinities better predict IgG IC binding to effector cells, suggesting that they may also help improve the estimation of *in vivo* cell response.

Next, we used the multivalent binding model with regression to predict *in vivo* antibody effector cell-driven platelet depletion in humanized mice. In the process of extending our previous model, we elected to use the cumulative density function of the exponential distribution as the link function in our generalized linear regression model to link the overall cell activity to the amount of target (e.g., platelet) depletion (Fig. 6a). Since the cell depletion effects have a limited range—one cannot deplete an antibody target of more than 100% or less than 0%—we must use a non-linear link function to transform the linear combination. While many functions provide this general relationship (such as the hyperbolic tangent function used before^16^), we realized that the extent of target cell depletion can be thought of as a form of survival analysis. In other words, given a certain antibody activity, a target cell has a certain probability of being cleared within the given timescale of the experiment. Assuming all target cells have an equal propensity of being cleared dictates an exponential relationship for the link function^21^.

**Figure 6:**
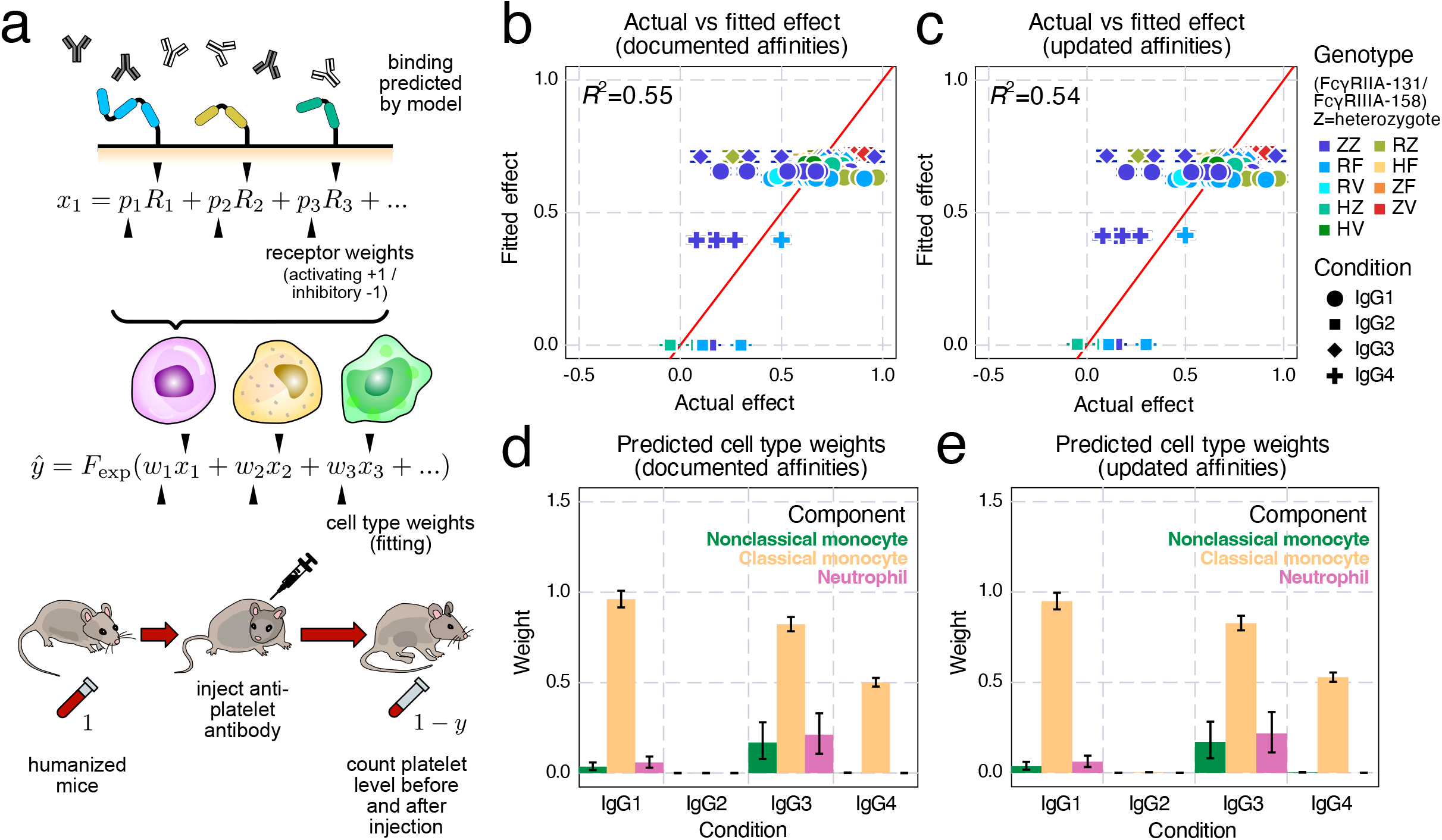
Perform *in vivo* target cell depletion regression on humanized mice. a) Schematic of *in vivo* platelet depletion regression. To predict the percentage decrease of platelet abundance after antibody injection in mice, we combined the binding model predictions with Fc receptor and effector cell type weights, then transformed the sum into depletion percentage with an exponential distribution cumulative density function. b-e) Results of regression ran on documented (b,d) and updated (c,e) affinities: b,c) Actual vs. predicted depletion of platelet; d,e) Predicted effector cell type weights; Error bars indicate the interquartile range from MCMC runs.

Having refined the cell clearance model, we applied it to a previously-collected dataset examining platelet depletion in humanized mice^22^. After fitting the cell type weighting, we found the model fit the experiments well, especially considering the experiment-to-experiment variability due to donor graft variation and other sources of experimental uncertainty (Fig. 6b/c). The fitting was almost identical when using documented (Fig. 6b) or updated (Fig. 6c) FcγR affinities.

A benefit of the generalized linear regression model is that it provides an easy interpretation of each component. Inspecting the cell type weights, we found that classical monocytes were inferred to be the predominant effector cell type (Fig. 6d/e). IgG2 had some binding to each effector cell type, but no activity was inferred whatsoever (Fig. 5f, 6d/e). As the affinity updates are most relevant to IgG2, and this isotype had no *in vivo* effect, it is reasonable that these changes had little effect on agreement with the data (Fig. 5f, 6e). While classical monocytes were inferred to exert the greatest impact on platelet depletion across isotypes, neutrophils, not classical monocytes, had the greatest binding (Fig. 5a–d). This demonstrates that the most bound cell type does not equate to the most potent effector. The regression model can incorporate the molecular level binding estimation and the depletion outcome to provide insights into the overall potency of each cell type. Overall, we found that the binding model could predict antibody-elicited effector responses *in vivo* in humanized mice.

## Discussion

In this work, we explored the binding properties of ICs with mixed IgG Fc composition and linked their *in vitro* effects to *in vivo* effector cell-elicited platelet depletion. To quantify the binding of mixed IgG ICs *in vitro,* we measured every human IgG subclass pair across a range of compositions multimerized at two different valencies (Fig. 1). Fitting these measurements to a model of multivalent interactions using documented affinities for each interaction, our model captured overall binding trends, with some outliers (Fig. 3). We uncovered that the model discrepancies could be explained by inaccurate estimates of especially low-affinity Fc receptor interactions, most prominently involving IgG2. We validated revised affinities within an independent dataset and found it greatly improved concordance with the data there as well. Finally, we used measurements of binding to effector cell populations to predict *in vivo* antibody-driven depletion of platelets in humanized mice (Fig. 5 & 6). While the updated affinities did not change the agreement of the model with the data, it did change the interpretation of IgG2’s small effect on depletion—rather than not binding to classical monocytes, IgG2 binds strongly when in a larger IC, but platelets might provide insufficient avidity to observe sufficient engagement (Fig. 6).

Considering that polysaccharide antigens present during bacterial infections or upon vaccination efficiently trigger IgG2 responses^23^, our data would support the notion that FcvR-dependent effector functions such as phagocytosis of opsonized bacteria may contribute to protective IgG responses in humans more than expected. Reversely, autoreactive IgG2 responses observed during many autoimmune diseases may contribute to autoimmune pathology via FcγRs, which may warrant to develop therapeutic interventions blocking this pathway also in IgG2-dominated autoimmune diseases^24^. Finally, with respect to the use of human IgG2 antibody formats as immunomodulatory antibodies for the therapy of cancer, our results would support strategies to engineer IgG2 variants with reduced binding to activating FcγRs and optimized binding for the inhibitory FcγRIIb, which has been shown to be critical for immunomodulatory IgG activity to further improve their therapeutic activity and reduce unwanted side-effects^25^.

IgG subclasses and glycan variants are defined by their differing affinity toward each Fc receptor^3,19,26^. Therefore, accurate measurements of each Fc receptor affinity are critical to understanding the differences in immune responses to each IgG. Using a mechanistic multivalent binding model alongside *in vitro* binding fluorescence measurements, we were able to derive a new set of Fc affinities distinct from those measured by surface plasmon resonance (SPR). Due to the heightened avidity, we found that multivalent ICs are better at detecting low-affinity IgG-Fc receptor interactions (Fig. 4b). Examining binding through ICs also better simulates the relevant structure of Fc-FcR interactions *in vivo.* Harnessing avidity to overcome the low affinity of interactions is a common theme in immunology and its experimental characterization. For instance, tetramers are routinely used for isolating antigen-selective T cells^27^. Here, we additionally show that these complexes can be used alongside quantitative models to infer properties of these systems.

Our results suggest that, within ICs comprised of several distinct subclasses or glycosylation variants, the Fc interaction effects are a blend of the constituent species’ properties. This means that ICs’ most extreme binding and effector responses should predominantly arise from whichever species is most potent in soliciting binding or a response. It also should provide some encouragement that the effector responses elicited from therapeutic antibodies should vary roughly in proportion to their relative composition, and small contaminants of alternative Fc subclasses or glycosylation can only have a substantial effect if those species differ extremely in their responses alone. One caveat of this observation is that we only examined mixtures of antibodies with differing Fcs but identical antigen binding—polyclonal mixtures of antibodies will have still other interaction effects, in part because antigens can form a higher valency complex when they are present in combination^28^. While in this work we only demonstrated Fc subclasses mixtures, the same lessons likely apply to glycosylation mixtures, both *in vitro* and *in vivo,* since different subclasses and glycosylation variants exert their effect through divergent affinities toward Fc receptors.

Fc receptor-mediated effects are central to protection from both endogenously produced and therapeutic antibodies. Our work demonstrates that computational methods greatly facilitate reasoning about the complex signaling of the Fcγ receptor pathway quantitatively and at both cellular and organismal levels. This work extends our previous modeling to the depletion of platelets as a target and humanized mice^16^. Humanized mice serve as an ideal surrogate for understanding human immunity^29^. However, this model system is complicated by graft-to-graft differences, including the level of humanization and genetic heterogeneity of human stem cell donors^29^. The depletion data reflected these complications, with high donor-to-donor and mouse-to-mouse variation^22^ (Fig. 6b/c). We anticipate that mechanistic models of antibody-mediated protection, such as the one here, will continue to grow in their utility for studying model systems such as humanized mice. In fact, as other features of antibodies are incorporated, such as variation in antigen specificity, it may become possible to connect behavior *in vitro* all the way to protection in human subjects^30,31^.

## Methods

### Data and Software Availability

All analysis was implemented in Julia and can be found at https://github.com/meyer-lab/FcRegression.jl.

### Chinese hamster ovary (CHO) cell FcγR Expression Quantitation

Human FcyR expression on stably transfected CHO cells was quantified by determining the antibody binding capacity (ABC) for antibodies specific for the respective Fcγ receptor (Tbl. S2)^13^. Quantum Simply Cellular (QSC) anti-mouse beads (Bangs Laboratories Ltd.) with known binding capacities for mouse IgG were used according to manufacturer’s instructions. Subsequently, a reference curve was generated by correlating the fluorescence intensity (caused by the respective anti-FcγR antibody) and the number of antibody binding sites of the different QSC beads. This reference curve was established in each experiment for all FcγR-specific antibodies of interest (PE-conjugated clone 10.1 to detect FcγRI, clone AT10 to detect FcγRIIa/b and clone 3G8 to detect FcγRIIIa, all from Biolegend) and used to calculate receptor numbers based on fluorescence intensity of FcyR staining on CHO cells. Samples were measured on a FACSCantoII flow cytometer and analyzed with FACSDiva software.

### Generalized multi-ligand, multi-receptor multivalent binding model

To model polyclonal antibody-antigen immune complexes (ICs), we employed a multivalent binding model to account for ICs of mixed IgG composition previously developed and detailed in Tan and Meyer^20^.

In this model, we define *N_L_* as the number of distinct monomer Fc’s and *N_R_* as the number of FcRs, and the association constant of monovalent Fc-FcR binding between Fc *i* and FcR *j* as *K_a,ij_*. Multivalent binding interactions after the initial interaction are assumed to have an association constant of 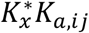, proportional to their corresponding monovalent affinity. The concentration of complexes is *L*_0_, and the complexes consist of random ligand monomer assortments according to their relative proportion. The proportion of ligand *i* among all monomers is *C_i_*. By this setup, we know 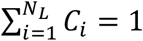. *R_tot,i_* is the total number of receptors *i* expressed on the cell surface (where this term is used synonymously for the actual determined number of binding sites for the respective anti-FcR antibodies), and *R*_eq,*i*_ the number of unbound receptors *i* on a cell at the equilibrium state during the ligand complex-receptor interaction.

The binding configuration at the equilibrium state between an individual complex and a cell expressing various receptors can be described as a vector **q** = (*q*_10_, *q*_11_,…, *q*_1*N_R_*_, *q*_20_,…,*q*_2*N_R_*_,*q*_30_,…,*q_N_L_N_R__*) of length *N*(*N_R_* + 1), where *q_ij_* is the number of ligand *i* bound to receptor *j*, and *q*_10_ is the number of unbound ligand *i* on that complex in this configuration. The sum of elements in **q** is equal to *f*, the effective avidity. For all *i* in {1,2,…,*N_L_*}, let 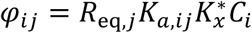 when *j* is in {1,2,…,*N_R_*}, and *φ*_10_ = *C_i_*. Then, the relative number of complexes in the configuration described by **q** at equilibrium is

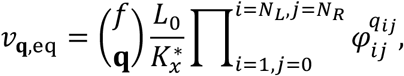

with 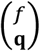 being the multinomial coefficient. Then the total relative amount of bound receptor type *n* at equilibrium is

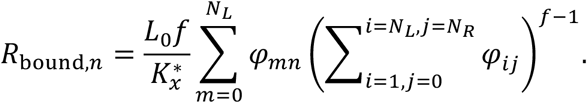

By conservation of mass, we know that *R_tot,n_* = *R*_eq,*n*_ + *R*_bound,*n*_ for each receptor type *n,* while *R*_bound,*n*_ is a function of *R*_eq,*n*_. Therefore, each *R*_eq,*n*_ can be solved numerically from its *R_tot,n_* measured experimentally. Similarly, the total relative number of complexes bind to at least one receptor on the cell is

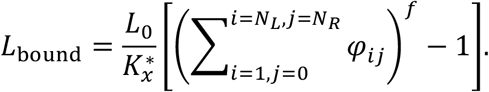

FcRs are activated through crosslinking. The amount of each kind of receptor in a multimerized complex can be calculated as

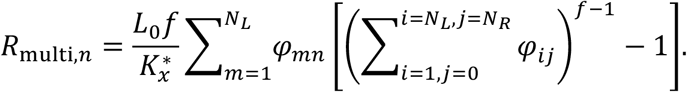

### Immune Complex Binding Measurement

Chinese hamster ovary (CHO) cells stably expressing human FcγRs were used to assess IgG-IC binding to hFcγRs as previously described^6^. Briefly, ICs were generated by coincubation of 10 μg/ml anti-TNP human IgG subclasses (clone 7B4) and 5 μg/ml BSA coupled with either an average of 4 or 33 TNP molecules (Biosearch Technologies) to mimic low or high valency ICs, respectively, for 3 h with gentle shaking at room temperature. To address the impact of distinct subclass combinations on hFcγR binding, human IgG1 through IgG4 subclasses were mixed at specific conditions (100%, 90%, 66%, 33%, 10% of one subclass filled up to 10μg/ml with the respective second subclass) before the addition of TNP-BSA. ICs were subsequently incubated with 100,000 FcγR expressing or untransfected control CHO cells for 1 h under gentle shaking at 4°C. Bound ICs were detected using a PE-conjugated goat anti-human IgG F(ab’)2 fragment at 0.5 μg/ml (Jackson ImmunoResearch Laboratories) on a BD FACSCanto II flow cytometer. Each IC binding measurement was normalized to the average of all the points within that replicate.

Alternatively, binding to human primary peripheral blood leukocytes co-expressing specific FcγRs was studied. Blood was drawn from healthy volunteers with informed consent of the donor and the local ethical committee. Erythrocytes were lysed by the addition of ddH_2_O for 30 sec at room temperature to obtain total leukocytes. Immune complexes were generated as described above and incubated with 200,000 leukocytes. Leukocyte subpopulations were identified by staining of cell-type-specific surface markers. Fluorescently labeled antibodies PE/Cy7-conjugated anti-CD19, PerCP-conjugated anti-CD3, APC-conjugated anti-CD33, Brilliant Violet 510 conjugated anti-CD14, FITC-conjugated anti-CD56 and APC-Fire 750 conjugated anti-CD45 were obtained from Biolegend.

The cell identification strategy was as follows: aggregates of cells were excluded by their forward light scatter (FSC) characteristics (area vs. height) and dead cells based on staining with DAPI. Leukocytes were identified by expression of common leukocyte marker CD45. Among those, neutrophils were gated based on high side light scatter (SSC) characteristics and lack of surface CD14 and classical monocytes based on intermediate SSC and expression of CD14. Within the CD14-SSC^low^ cells, B and T cells were gated by expression of CD19 or CD3, respectively. Staining of CD56 was used to distinguish NK cells. The remaining CD33 expressing cells are gated as non-classical monocytes.

Data were analyzed with FlowJo or FACSDiva Flow Cytometry Analysis Software. The relative fluorescence units were normalized so that measurements of each day have geometric means of 1.0. The variance explained calculated for principal component analysis was defined as 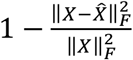, where 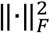 indicates the Frobenius norm.

### Immune Complex Binding Analysis

Fitting the parameters in the binding quantification was performed by Monte Carlo Markov Chain (MCMC) implemented by Turing.jl^32^.

At first, we plugged in the documented values into the binding model for all parameters without fitting, thus the geometric means of CHO cell receptor expression (Tbl. S2), documented affinities^3^, nominal valencies (4 and 33), and 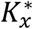 as 6.31 × 10^-13^cell · M^16^, as estimated in previous work (Fig. 3c)^33^. To examine the role of affinity fitting, we used MCMC to fit all parameters except (Fig. 3d) and including (Fig. 3e) affinities. CHO receptor prior distributions were inferred from their measured values through maximal likelihood estimation (MLE) in Distributions.jl^34^ for both IgG mixture dataset (Tbl. S2) and validation Robinett et al.^33^ dataset (Tbl. S5). The affinity priors were inferred from documented Fc affinities and standard errors with these rules: (1) each prior follows a lognormal distribution; (2) the mode of the distribution is the documented value, and the interquartile range of the distribution is the standard error; (3) if the values of mode or standard errors are too small, the mode was clipped to 1 × 10^4^ M^-1^, and the interquartile range was clipped to 1 × 10^4^M^-1^ to deal with recorded nonbinding cases^3,35^. The priors of valencies and crosslinking constant were as follows

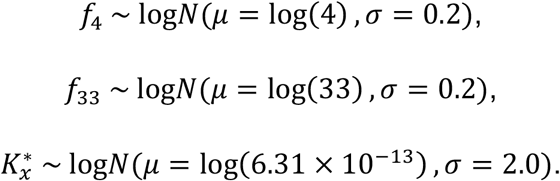

MCMC was initialized with the maximum a posteriori estimation (MAP) optimized by a limited-memory BFGS algorithm implemented by Optim.jl^36^, then sampled through a No U-Turn Sampler (NUTS) implemented by Turing.jl^32^.

### *In vivo* Regression Model

We extended the *in vivo* antibody-elicited target cell depletion regression model with both cell type weights and FcγR weights (Fig. 6a). Depletion, *y*, was represented as the percent reduction in the number of target cells.

To quantify the activity of each effector cell, we first used the multivalent binding model to predict the amount of multimerized FcγR of each kind, *R*_multi,*i*_, assuming each IC is 4-valent. Then the activity of this cell type is assumed to be a linear combination of these predictions and a set of cell type weights, *p_i_*, that are set to either +1 or −1 for activating or inhibitory receptors, respectively, clipped to 0 if it is negative:

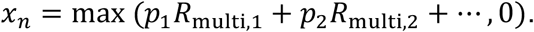

To determine how these cell types bring the depletion effect at the organism level, we combine their estimated effects, *x_n_*, with a weighted sum, where we introduce another set of weights, *w_n_*, that are specific to each cell type. To convert the activities to a limited range of depletion, (i.e., one cannot have a reduction over 100%), the regression was transformed by an exponential linker function (the cumulative density function of exponential distribution) such that the predicted effectiveness: *ŷ* = *F*_exp_(*wx*) = 1 - exp(-*wx*;) so that lim_*X*→∞_*F*_exp_(*X*) = 1. Together, we have the estimated depletion as

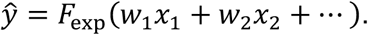

We did not estimate the amount of each cell type in an individual, nor did we include them in the model, because the weights, *w_n_*, were supposed to absorb these quantities, while requiring effector cell abundance can limit the application of this model to specific organs where the tissue resident cell abundance cannot be accurately quantified.

The regression against *in vivo* effectiveness of IgG treatments was performed via Monte Carlo Markov Chain (MCMC) implemented by Turing.jl^32^. For the multivalent binding model, the ligand concentration was assumed to be 1 nM and valency to be 4. The receptor expression level was set to the geometric means of the values measured in previous work (Tbl. S6)^13^. For the receptor weights, *p_i_*, we set the weight of the only inhibitory receptor, FcγRIIB, as −1.0, while every activating receptor to be +1.0.

MCMC was initialized with the maximum a posteriori estimation (MAP) optimized by a limited-memory BFGS algorithm implemented by Optim.jl^36^, then sampled through a No U-Turn Sampler (NUTS) implemented by Turing.jl^32^.

## Acknowledgements

This work was supported by NIH U01-AI-148119 to A.S.M. and F.N.

## Author contributions statement

A.S.M. and F.N. conceived the study. A.L., P.V., and M.B. performed the experiments. A.S.M., S.L., and Z.C.T. performed the computational analysis. All authors reviewed the manuscript.

