## Supplemental Figures & Tables for "Mixed IgG Fc immune complexes exhibit blended binding profiles and refine FcR affinity estimates"

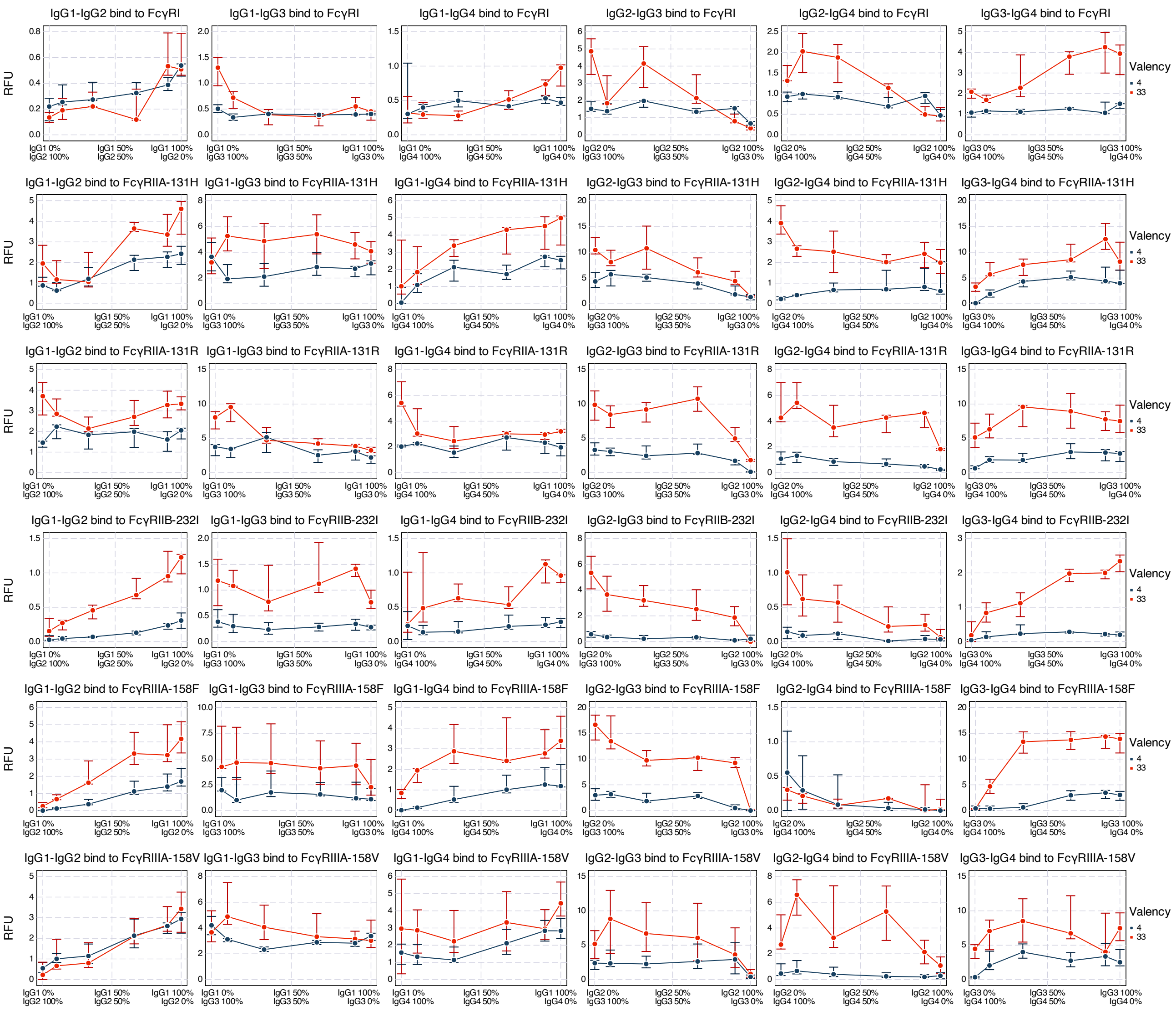


**Figure S1: Experimental IC mixture binding data.** Quantification of human IgG subclass pairs TNP-4-BSA and TNP-33-BSA IC binding to CHO cells expressing the indicated hFcγRs. Relative fluorescent units (RFU) of different multivalent immune complexes consisting of various IgG mixtures binding to different human immune cell receptors. Error bars indicate the biological replicates from experiments. Fluorescent values were normalized so that the daily geometric average measurements are 1.

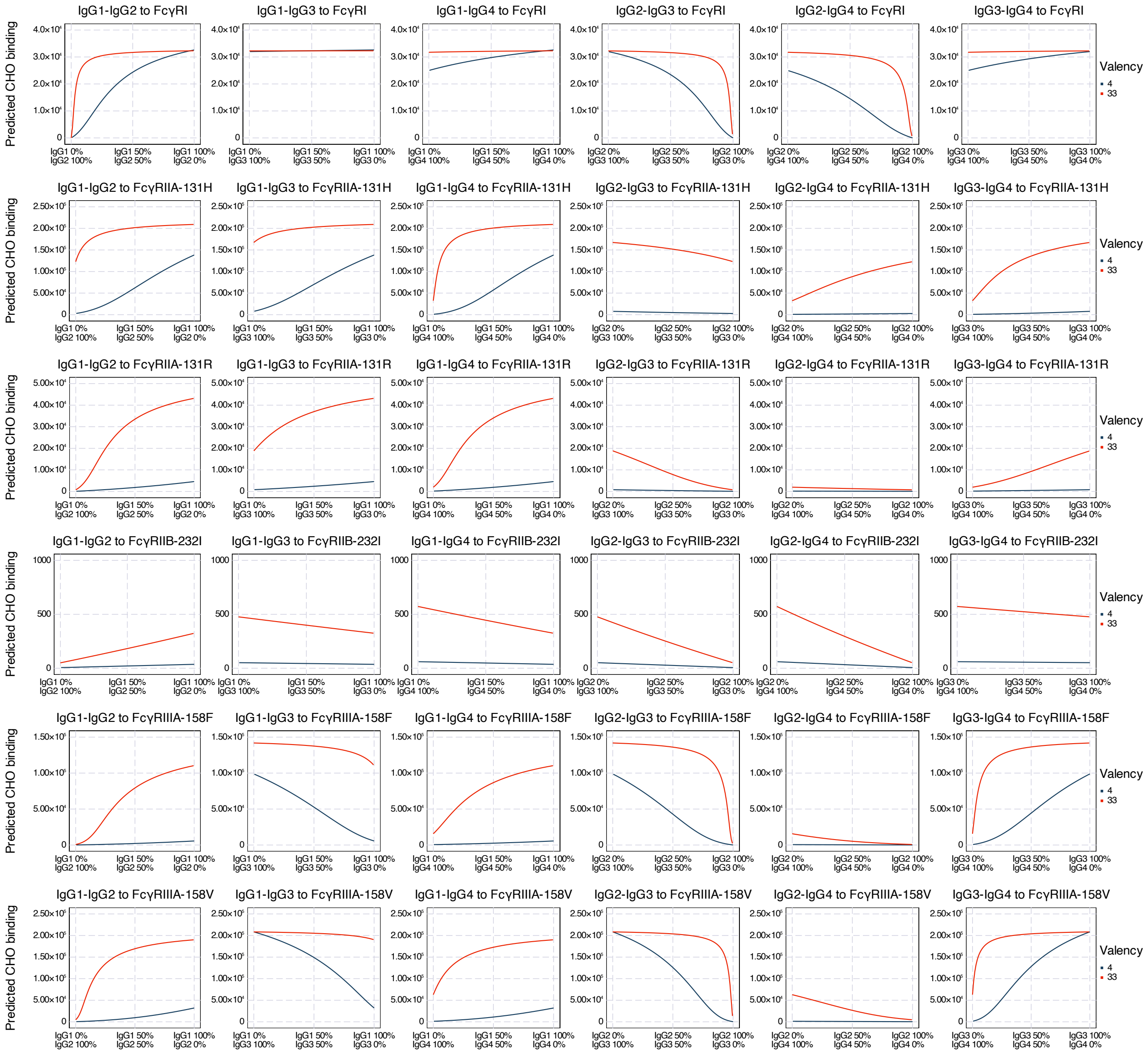


**Figure S2: Predicted binding of IgG subclass mixtures with documented affinities.** Amount of binding for complexes of each IgG subclass pairs binding to each CHO cell predicted by the multivalent binding model with the documented affinities. The receptor abundances were as geometric means of measurement.

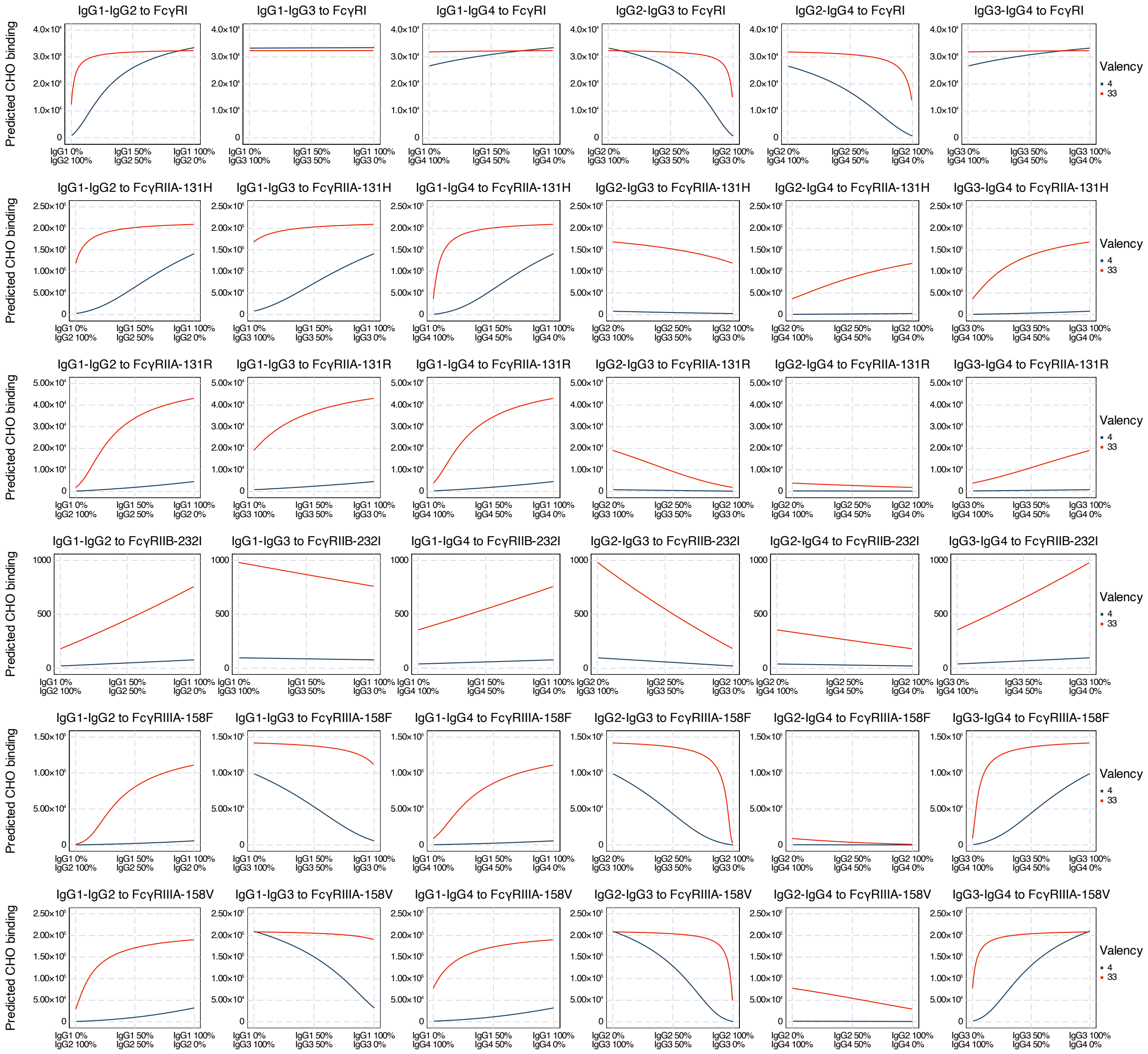


**Figure S3: Predicted binding of IgG subclass mixtures with updated affinities.** Amount of binding for complexes of each IgG subclass pairs binding to each CHO cell predicted by the multivalent binding model with the updated affinities. The receptor abundances were as geometric means of measurement.

### Supplemental Tables

Table S1. One-way ANOVA performed on the measurements indicates that the majority of variance in measurements comes from between conditions.

| Source | DF | SS | MSS | F | p |
|---|---|---|---|---|---|
| Condition | 431 | 11110.23 | 25.778 | 6.00915 | $5.9926\times10^{-125}$ |
| Residuals | 1066 | 4572.88 | 4.290 | | |
| Total | 1497 | 15683.11 | | | |

$R^2$ = 0.7084

Table S2. Geometric mean and inferred prior distribution of FcR abundance. Antibody binding capacity on CHO cells from measurements used in IgG mixture *in vitro* binding experiment. The geometric means were calculated from primary data published in previous work[1]. logN represents a lognormal distribution.

| Receptor | Geometric mean | Inferred distribution |
|---|---|---|
| FcγRI | 101494 | logN(μ=11.53, σ=0.26) |
| FcγRIIA-131H | 1006300 | logN(μ=13.82, σ=0.27) |
| FcγRIIA-131R | 190433 | logN(μ=12.16, σ=1.46) |
| FcγRIIB-232I | 75085 | logN(μ=11.23, σ=1.40) |
| FcγRIIIA-158F | 634324 | logN(μ=13.36, σ=0.70) |
| FcγRIIIA-158V | 979452 | logN(μ=13.80, σ=0.38) |

Table S3. Updated Fc affinities' interquartile ranges from their posterior distributions.

| $K_a$ ($M^{-1}$) | IgG1 | IgG2 | IgG3 | IgG4 |
|---|---|---|---|---|
| FcγRI | $5.809\sim8.635\times10^7$ | $1.189\sim1.911\times10^6$ | $5.525\sim8.730\times10^7$ | $3.012\sim4.770\times10^7$ |
| FcγRIIA-131H | $4.780\sim5.929\times10^6$ | $3.930\sim4.724\times10^5$ | $8.622\sim9.621\times10^5$ | $1.625\sim2.057\times10^5$ |
| FcγRIIA-131R | $3.277\sim3.758\times10^6$ | $1.719\sim2.339\times10^5$ | $8.521\sim9.861\times10^5$ | $2.833\sim3.864\times10^5$ |
| FcγRIIB-232I | $2.166\sim3.053\times10^5$ | $5.931\sim8.134\times10^4$ | $2.714\sim3.702\times10^5$ | $1.131\sim1.504\times10^5$ |
| FcγRIIIA-158F | $1.106\sim1.288\times10^6$ | $2.938\sim4.405\times10^4$ | $7.243\sim8.329\times10^6$ | $1.337\sim1.764\times10^5$ |
| FcγRIIIA-158V | $1.913\sim2.107\times10^6$ | $1.433\sim1.966\times10^5$ | $0.919\sim1.073\times10^7$ | $2.509\sim3.348\times10^5$ |

Table S4. Updated Fc affinities median values.

| $K_a$ ($M^{-1}$) | IgG1 | IgG2 | IgG3 | IgG4 |
|---|---|---|---|---|
| FcγRI | $7.140\times10^7$ | $1.496\times10^6$ | $6.998\times10^7$ | $3.835\times10^7$ |
| FcγRIIA-131H | $5.314\times10^6$ | $4.306\times10^5$ | $9.128\times10^5$ | $1.808\times10^5$ |
| FcγRIIA-131R | $3.503\times10^6$ | $2.005\times10^5$ | $9.178\times10^5$ | $3.291\times10^5$ |
| FcγRIIB-232I | $2.549\times10^5$ | $6.911\times10^4$ | $3.157\times10^5$ | $1.305\times10^5$ |
| FcγRIIIA-158F | $1.195\times10^6$ | $3.639\times10^4$ | $7.741\times10^6$ | $1.536\times10^5$ |
| FcγRIIIA-158V | $2.006\times10^6$ | $1.678\times10^5$ | $9.941\times10^6$ | $2.906\times10^5$ |

Table S5. Geometric mean and inferred prior distribution of FcR abundance. Antibody binding capacity on CHO cells used in validation with Robinett et al.[2] binding dataset. The geometric means were calculated from primary data published in previous work[1]. logN represents a lognormal distribution.

| Receptor | Geometric mean | Inferred distribution |
|---|---|---|
| FcγRI | 232872 | logN(μ=12.36, σ=0.25) |
| FcγRIIA-131H | 318819 | logN(μ=12.67, σ=0.26) |
| FcγRIIA-131R | 1605372 | logN(μ=14.29, σ=0.14) |
| FcγRIIB-232I | 394556 | logN(μ=12.89, σ=0.45) |
| FcγRIIIA-158F | 4677645 | logN(μ=15.36, σ=0.22) |
| FcγRIIIA-158V | 3680708 | logN(μ=15.12, σ=0.25) |

Table S6. Geometric means of measured FcγR expression, i.e. the number of quantified binding sites for the respective anti-FcR antibodies on effector cells calculated from the primary data published in previous work[1].

| Receptor | Non-classical monocyte | Classical monocyte | Neutrophil |
|---|---|---|---|
| FcγRI | 6326 | 84559 | 1847 |
| FcγRIIA | 82542 | 96646 | 158228 |
| FcγRIIB | 7140 | 5167 | 2351 |
| FcγRIIIA | 200213 | 19533 | 0* |
| FcγRIIIB | 0* | 0* | 1299166 |

* Although one cannot have a geometric mean as 0, these values were consistently measured as non-expressed, so we used 0 as the value.
